# Lubricin’s Mucin Domain Has Strong Polyproline Type-II Helical Character

**DOI:** 10.1101/2025.06.15.659778

**Authors:** Bibo ’Noah’ Feng, Ava J. Marks, Faith Kim, Mandar Naik, Tannin A. Schmidt, Khaled A. Elsaid, Gregory D. Jay, Brenda Rubenstein

## Abstract

Lubricin is a glycoprotein that is crucial for maintaining joint health by preventing joint wear by reducing joint friction in the boundary mode. Lubricin was recently observed to hinder the formation of uric acid crystals in the joint and prevent a form of gouty arthritis. However, despite lubricin’s great physiological importance, our current understanding of the molecular origins of lubricin’s beneficial properties is limited by a lack of detailed structural information regarding its central mucin domain: lubricin’s large size (227.5 kDa) and numerous glycosylations pose a substantial obstacle to conventional experimental methods for solving protein structures. In this work, we employ a combination of physics-based replica exchange molecular dynamics (REMD) simulations and circular dichroism (CD) experiments to shed light on the structure of lubricin’s central mucin domain. Using REMD, we model [KEPAPTTP]_2_, an amino acid repeat found throughout the mucin domain, and find that the mucin domain is likely to exhibit polyproline type II (PPII) helices, which are further stabilized by O-linked oligosaccharide chains. Motivated by these simulation results, we performed circular dichroism spectroscopy on fragments of the mucin domain that also show clear polyproline-II helical character, corroborating our computational findings. Altogether, this work provides strong evidence of a lubricin mucin domain with significant polyproline type II content. As polyproline helices are often also found in other glycoproteins with antifreeze properties, this work may also explain the atomistic underpinnings of their interfacial functions, including lubrication and competition with crystal formation.

**SIGNIFICANCE:** Lubricin is a mucinous glycoprotein containing a central heavily glycosylated domain that plays a crucial role in the lubrication of joints. However, little is known about the structure of this large central mucin domain, which makes the development of related therapeutics or biomedical devices challenging. In this work, we provide strong evidence that lubricin’s mucin domain possesses polyproline type II (PPII) character that is enhanced by glycosylation upon the basis of a combination of molecular dynamics simulations and circular dichroism experiments. Lubricin’s PPII character provides a molecular basis for its lubricating properties, which may provide insights into related antifreeze glycoproteins (AFGP) and the development of new biocompatible lubricants and ryo-preservatives.

## INTRODUCTION

Lubricin, also known as proteoglycan 4, is a glycoprotein deeply involved in promoting joint health in humans (see Figure 1) (1–3). It is a multi-domain protein that consists of an N-terminal domain, a C-terminal domain, and a central mucin domain composed primarily of KEPAPTTP tandem repeats (with occasional variations in the repeating unit) (see Figure 2). Heavily glycosylated with O-linked Gal NAc-Gal oligosaccharide chains, the central mucin domain has been implicated as the main driver behind many of lubricin’s beneficial properties, such as its ability to reduce wear of cartilage by acting as a boundary lubricant and as an anti-adhesive in the joints (4–6). Recently, it was also discovered that the presence of lubricin inhibits crystal nucleation of uric acid, a main driver of gout (7); however, the molecular mechanisms behind such phenomena are still unclear.

**Figure 1:**
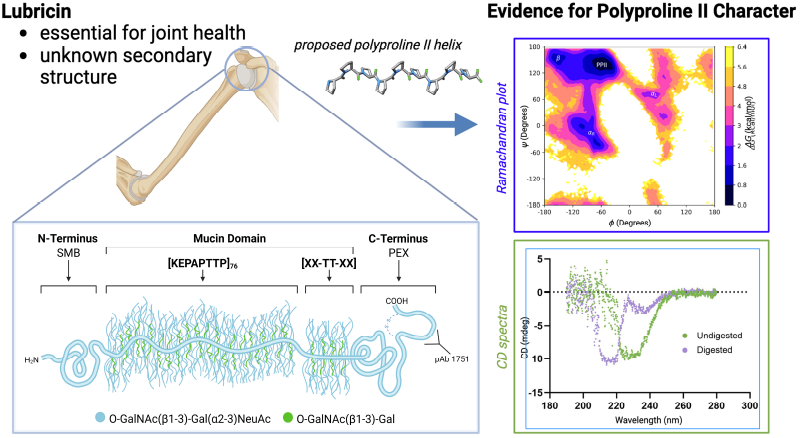
(Table of Contents Image) In this manuscript, we demonstrate that lubricin’s mucin domain possesses polyproline type-II helical character using a combination of molecular dynamics simulations and circular dichroism spectroscopy.

**Figure 2:**
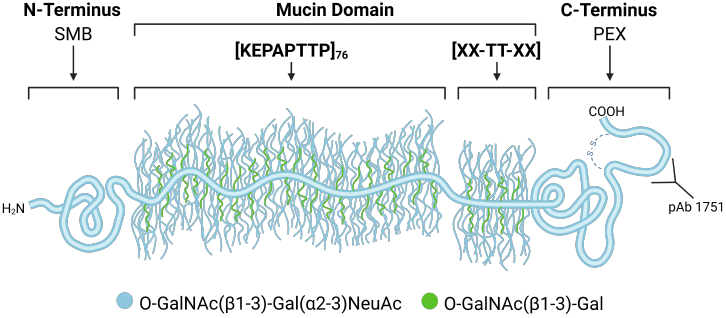
A schematic of lubricin, which consists of an N-terminal domain, a central mucin domain with many KEPAPTTP repeats, and a C-terminal domain referred to as a hemopexin (PEX) domain. The C-terminus possesses a disulfide-bonded peptide resulting from subtilisin cleavage (20). Common glycosidic side chains found on lubricin are also depicted.

Despite lubricin’s importance in maintaining joint health, our understanding of its mechanism of action is limited by a lack of detailed structural data regarding its central mucin domain. Due to lubricin’s large size (227.5 kDa) and numerous glycosylations, determining its structure via conventional experimental methods, such as X-ray crystallography and nuclear magnetic resonance (NMR), is very challenging. Furthermore, although recently developed deep learning-based methods such as AlphaFold have made major breakthroughs in accurately determining protein structures that are difficult to solve via experimental means, these methods currently struggle with heavily glycosylated proteins such as lubricin (8, 9).

Currently, the limited structural data available in the literature on lubricin’s mucin domain come from atomic force microscopy, which illustrates how lubricin’s mucin domain can self-assemble into a large helical structure when the globular end domains bind to a surface (5). Although atomic force microscopy imaging provides information on the general shape of the mucin domain when the terminal domains are bound to a surface, the structure observed in these images cannot be generalized as lubricin is not always bound to a surface (10, 11) and is, in fact, often found in a myriad of biological fluids such as tears (12) and in the amniotic membrane (13). As such, crucial atomistic structural details that could be relevant for understanding the mucin domain’s function are missing. One important but currently unknown piece of structural information is the secondary structure of the mucin domain. Knowledge of the domain’s secondary structure would provide information regarding the typical spacing between nearby glycan chains, which may in turn provide insights into how lubricin interacts with other molecular species, such as uric acid crystals. Hence, determining the secondary structure of the mucin domain may improve our understanding of lubricin’s mechanism of action. For example, recent non-equilibrium molecular dynamics simulations showed that the presence of O-linked glycosylations in the mucin domain contributes to Newtonian solution viscosity by interfering with mucin entanglements (14).

Despite the lack of experimental secondary structural data for the mucin domain, one can make a reasonable guess regarding the types of secondary structures possible in the domain based on its amino acid sequence and post-translational modifications, especially due to the domain’s repetitive nature. As the mucin domain is comprised mostly of KEPAPTTP tandem repeats, prolines are evidently the most common residue, making up approximately ∼31% of the mucin domain. Since prolines are widely interspersed throughout the mucin domain, the two most common secondary structures — *α*-helices and *β*-sheets — are unlikely to be found in the domain, as prolines are known to disrupt the formation of both of these secondary structures (15, 16). Although the absence of *α*-helices and *β*-sheets may sometimes indicate that a protein is intrinsically disordered, structural order at the secondary level can still be achieved via other, albeit less common, types of secondary structures. A class of protein secondary structures that could comprise the mucin domain are polyproline II (PPII) helices. The PPII helical conformation is characterized by backbone dihedral *ϕ* /*ψ* angles of −75^°^ /145^°^, resulting in a left-handed helix with 3 residues per turn (Figure 3) (17). Unlike *α*-helices and *β*-sheets, PPII helices neither possess regular intrachain nor interchain hydrogen bonding between backbone atoms. Instead, the formation of the PPII conformation is driven by backbone solvation and the reduction of unfavorable steric interactions (17, 18). Due to geometric constraints imposed by their sidechains, prolines possess the highest propensity to form PPII helices, but this conformation is accessible to other residues as well (18, 19). For instance, alanine and residues with long linear side chains, such as lysine, were demonstrated to also possess a high propensity for adopting a PPII helical conformation. Hence, five out of the eight total residues in a typical mucin domain tandem repeat (KEPAPTTP) possess high propensities for the PPII conformation.

**Figure 3:**
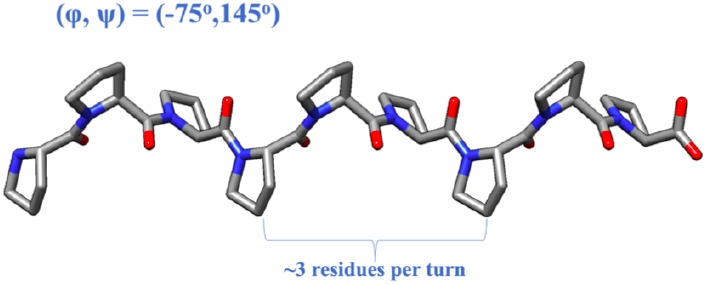
A schematic of a polyproline Type-II (PPII) helix. PPII helices are characterized by backbone dihedral *ϕ*/*ψ* angles of −75^°^/145^°^, which result in a left-handed helix with three residues per turn. PPII helices also possess limited hydrogen-bonding.

Beyond the amino acid composition in a peptide or protein, post-translational modifications such as glycosylations could also influence the formation of PPII helices. For instance, Rani *et al*. demonstrated that O-linked GlcNAcylation decreases the free energy of the PPII helical conformation for a model threonine dipeptide system (21). Another example is the antifreeze glycoprotein 8 (AFGP8): REMD has illustrated that threonine-linked GalNAc-Gal glycosylations — which, interestingly, are the same glycosylations in lubricin’s mucin domain — promote an organized and compact amphipathic PPII helical structure, allowing glycoproteins to inhibit ice recrystallization at high molecular weights (22, 23).

Given that lubricin’s mucin domain contains many characteristics that should promote the PPII conformation, in this work, we hypothesize that PPII helices are the dominant secondary structural elements present in the mucin domain. To analyze this hypothesis, we first utilized REMD to sample the conformational landscape of model peptides that are representative of the mucin domain, ultimately demonstrating that PPII helices are likely their most energetically favorable secondary structure. Follow-up simulations moreover demonstrate that the stability of the PPII conformation is even further increased upon glycosylation, which would be present as post-translational modifications. With this computational evidence as motivation, we then analyzed the PPII content of mucin domains using circular dichroism (CD) spectroscopy. These studies similarly point to PPII character, corroborating our computational predictions. Our results offer novel insights into the structure of lubricin’s mucin domain, which we hope can be used in the future to facilitate understanding of the mechanisms behind lubricin’s medically-beneficial properties (24, 25) and other related proteins’ cryogenic properties, and the development of more potent therapeutics.

## METHODS

### Computational Approach

To identify stable secondary structures within lubricin’s mucin domain, we used REMD simulations to calculate a Ramachandran-like free energy surface for our model mucin domain. We then compared this surface to those expected for a variety of alternative secondary structures to assess the likely secondary content of the domain.

### Model of the Mucin Domain

The full mucin domain of lubricin consists of hundreds of glycosylated residues, making it too large for direct simulation. To reduce the computational costs of our simulations of lubricin’s mucin domain, we make the following assumption: since the mucin domain’s amino acid sequence composition is highly repetitive, and since secondary structures are mostly local structures, any secondary structure which emerges in a small subsection of the mucin domain should be generalizable to the rest of the domain. Hence, instead of simulating the entire ∼ 500-residues-long mucin domain, we work under the assumption that simulations of model peptides composed of only a few tandem repeats within the mucin domain should still provide useful information regarding the domain’s secondary structural content.

In this work, we thus focus on an important structural subsection of the mucin domain corresponding to residues 393-410 (PKEPAPTTTKEPAPTTPK), which encompasses both the KEPAPTTP tandem repeat and the most common variation from the tandem repeat (KEPAPTTT). Both the glycosylated (glycopeptide) and non-glycosylated versions of the model peptide were built with the aim of understanding the effect of glycosylation on the peptide’s secondary structure. Initial non-glycosylated peptides were constructed using UCSF’s Chimera structure building tool (26) (Figure 4). (*ϕ, ψ*) dihedral angles of every residue were initially set to (−75^°^, 145^°^), the polyproline type II (PPII) helical conformation, as we expected PPII helices to be a common structural element in the mucin domain. While initializing the domains in this way has the potential to bias the final structures attained, as we will further discuss below, we find that this initialization does not inhibit related peptides from attaining alternative conformations and hence practically imposes a limited bias. For the glycopeptide, the CHARMM-GUI’s Glycan Modeler (27) was used to attach O-linked GalNac-Gal-glycans to threonines in the initial peptide generated using Chimera. Since residue Thr^401^of lubricin (corresponding to the third threonine in the model peptide) is not glycosylated (28), glycosylations were only attached to the other four threonines in the model peptide.

**Figure 4:**
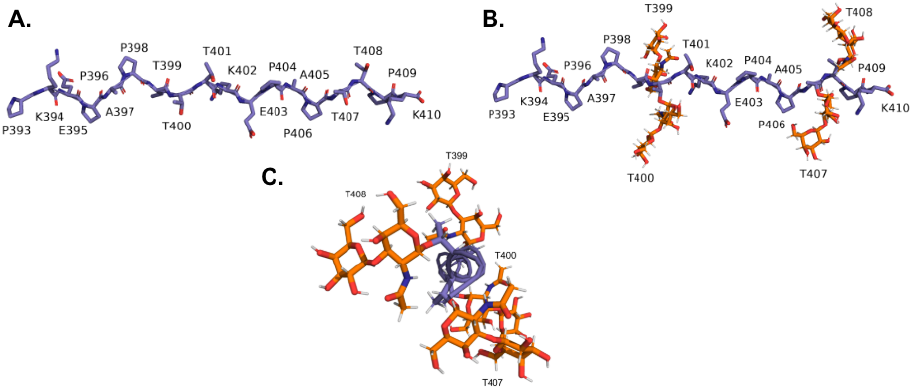
A) Non-glycosylated and B) glycosylated subsections of the mucin domain modeled in this work. These subsections correspond to residues 393-410 (PKEPAPTTTKEPAPTTPK), which contain both the KEPAPTTP and KEPAPTTT repeat units. C) End-on view of the glycosylated model peptide illustrating the positioning of the glycosylations around the peptide unit.

### Replica Exchange Molecular Dynamics Simulations

To further reduce the computational costs of and accelerate our modeling, we employ an enhanced sampling technique called REMD (29, 30). In REMD simulations, multiple MD simulations of a system called a replica are conducted in parallel at different temperatures. The temperatures of the replicas typically span a wide range, so the higher temperature replicas have enough thermal energy to overcome large energy barriers, while the lower temperature replicas are at temperatures of interest for the system. Exchanges between configurations sampled by neighboring temperature replicas are periodically made, which accelerates the sampling of the full energy landscape by allowing low-temperature simulations of interest to escape local minima. However, in order to ensure that the correct Boltzmann ensemble averages for a replica at any given temperature are preserved, exchanges between neighboring temperature replicas can only be made with a certain probability, which depends on the temperatures and the configurations being exchanged (31, 32).

REMD simulations were performed for both the glycosylated and non-glycosylated model peptides. All REMD simulations were conducted using GROMACS, a free, open-source software suite for high-performance MD simulations (33). Initial molecular topologies of the model peptides were generated via the CHARMM-GUI and the CHARMM36m force field (34). Simulations were conducted with the TIP3P water model (35) in an octahedral box extending 1 nanometer beyond the solute in all three dimensions. Before each production run, gradient-descent energy minimization was performed until the force on every atom was less than 1000 kJ/mol/nm. NVT (constant number of atoms, constant volume, and constant temperature) and NPT (constant number of atoms, constant pressure, and constant temperature) equilibration runs, both 200 ps in length with 2 fs timesteps, were then conducted with a restraining force constant of 400 kJ/mol/nm^2^ for the backbone atoms, 40 kJ/mol/nm^2^ for the sidechain atoms, and 4 kJ/mol/rad^2^ for the dihedral angles. After equilibration, 200 ns NPT production runs with 2 fs timesteps were performed without restraints. Each production run contains 64 replica trajectories, with temperatures distributed according to a geometric progression between 300 K and 455.95 K. Exchange attempts between neighboring temperature replicas were made every picosecond. To ensure equilibration, only the last 100 ns of each simulation was used for analysis.

### Free Energy Landscape Calculations

The enhanced sampling capabilities of REMD simulations result in a greater exploration of a protein’s free energy landscape, enabling greater insights into a protein’s structural characteristics. For instance, REMD simulations performed by Mochizuki *et al*. were used to illustrate that the global free energy minimum of AFGP8 lies within the PPII helical region of the Ramachandran plot (22). The present work will adopt a similar approach as that employed by Mochizuki *et al*. to examine the secondary structural free energy landscape for model peptides of lubricin’s mucin domain. For a given pair of dihedral angles (*ϕ, ψ*) on the Ramachandran plot, the free energy relative to the global free energy minimum can be calculated using:

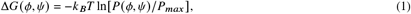

where *k* _*B*_ is Boltzmann’s constant, *T* is the temperature, *P* (*ϕ, ψ*) is the probability that an amino acid within the protein has dihedral angles (*ϕ, ψ*), and *P*_*max*_ is the probability of the most frequently occupied (*ϕ, ψ*)pair within the protein. The probability distribution, *P* (*ϕ, ψ*), can be estimated by partitioning dihedral angles sampled by an REMD simulation into 2D histogram bins that span the entire dihedral angle phase space. The free energy for each bin can then be calculated using equation (1), resulting in the construction of a free energy surface. The resulting free energy histogram can then be compared to that of known secondary structures to assess lubricin’s secondary structural content.

## Experimental Methods

### Recombinant Human PRG4 and Polyclonal Antibody Directed Towards the PRG4 C-Terminus

Recombinant human PRG4 (rhPRG4) (Lubris, LLC, Naples, FL) was expressed by CHO-M cells as previously described (36). CHO-M cells were transfected with the full-length human PRG4 gene resulting in a full-length PRG4 (1404 amino acids in length). The presence of O-linked glycosylations comprised of Gal-GalNAc (Core 1) was confirmed by mass spectrometry (37), which showed that the majority are sialyated (38). Polyclonal Antibody (pAb) 1751 (Figure 2) was manufactured against the PRG4 sequence Q^1279^KCPGRRPALNYPVYGETTQ^1298^, which is adjacent to the subtilisin cleavage site in the C-terminus of PRG4. This 20-mer peptide sequence was manufactured and conjugated to keyhole limpet antigen using standard methods and then injected into two New Zealand white rabbits (BioMatik, Wilmington, DE). Screening of rabbit sera for immunopositivity against rhPRG4 was conducted by ELISA.

### Partial Digestion of rhPRG4

Recombinant human proteoglycan-4 (rhPRG4) (Lubris, Naples, FL) was digested using cathepsin G (Sigma-Aldrich, Saint Louis, MO, CAS 107200-92-0) at a protease:protein weight-to-weight ratio of 1:500 for 1 hour at 37°C with shaking (39). After 1 hour, the proteolytic inhibitors aprotinin (Sigma-Aldrich, Saint Louis, MO, CAS 9087-70-1) and leupeptin (Sigma-Aldrich, Saint Louis, MO, CAS 147385-61-3) were added at a 1:1 concentration ratio. The digestate was fractionated using Fast Protein Liquid Chromatography (FPLC) on a HiPrep Sephracryl S-500 HR column (Cytiva, Marlborough, MA) using the ÄKTA Pure™ chromatography system (Cytiva, Marlborough, MA) at 0.75 ml/min in 1M NaCl in PBS at pH 7.4. Eluted fractions were collected based on their absorbance peaks at 280 nm. Collected fractions were dialyzed overnight in Spectra/Por™ 3 RC Dialysis Membrane Tubing (3500 Da MWCO) (Spectrum Laboratories Inc., Rancho Dominguez, CA) against phosphate buffed saline (PBS) at 4°C. Samples were then concentrated to a final volume of 500 µL using Amicon® Ultra Centrifugal Filters (10 kDa MWCO) (Merck KGaA, Darmstadt, Germany) and analyzed via sodium dodecyl sulfate acrylamide gel electrophoresis (SDS-PAGE) on a NuPAGE™ 4-12%, Bis-Tris, 1.0-1.5 mm, Mini Protein Gel (Invitrogen, Waltham, MA) and stained with GelCode Blue (Thermo Fisher Scientific Inc., Waltham, MA). Additional gels were Western blotted and probed with the pAb 1751 antibody and secondary antibody goat anti-rabbit IgG horseraddish peroxidase (Invitrogen, Waltham, MA). The apparent highest molecular weight samples from three separate digestions were pooled together based on the FPLC retention time, SDS-PAGE, and Western blot results. This pooled sample was exhaustively dialyzed a second time against distilled deionized (DDI) water for circular dichromatism (CD) spectroscopy.

### Circular Dichroism Spectroscopy

Partially Cathepsin-G digested rhPRG4 was analyzed by circular dichroismm (CD) spectroscopy (40) using a J-815 Spectropolarimeter (Jasco Inc., Easton, MD), coupled with a PTC-423S temperature controller (Jasco Inc., Easton, MD) and a Peltier controlled water bath (Jasco Inc., Easton, MD) at 20°C. A volume of 600 µL of the digested rhPRG4 was pipetted into a quartz cuvette where DDI water was used as the baseline measurement. The spectral range was 190-280 nm with a data pitch of 0.1 nm. The digital integration time (DIT) was set to 1 second, and the spectral bandwidth was set to 1 nm. Scanning speed was fixed at 20 nm/min, and the spectra were measured in triplicate. Analysis of the CD data was performed using the Spectra Manager software (Jasco Inc., Easton, MD) and the BeStSel algorithm which is designed to provide detailed secondary structure information including the content of PPII helices, beta-sheets, and disordered regions (41).

## RESULTS

### Convergence Checks for REMD Simulations of Model Lubricin Peptides

The accuracy of the results obtained with molecular dynamics simulations depends greatly on the quality of their sampling. To verify that sufficient sampling has been achieved for a simulation, one can compare the distributions of some physical property sampled during two independent time intervals. Significant overlap between the two distributions likely indicates high convergence and adequate sampling. A common choice of a physical variable used for convergence checks is the radius of gyration *R*_*g*_ of a protein, a quantitative measure of a protein’s compactness that is defined by:

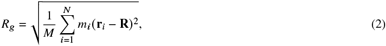

where *M* is the total mass, **R** is the center of mass, *N* is the total number of atoms, and *m*_*i*_ and **r**_*i*_ are the mass and position of the *i*^*th*^ atom, respectively.

Hence, to check the convergence of our REMD simulations, the radius of gyration *R*_*g*_ for the lubricin model peptide with and without glycosylations was calculated at every picosecond in the 300 K replica trajectories (the replica at the temperature of interest on which further analysis will be performed). *R*_*g*_ probability distributions were then estimated using histograms with 50 bins for two independent time intervals of the non-glycosylated peptide and the glycopeptide trajectories at 300 K. High degrees of overlap (*>* 90% overlap) are observed between the *R*_*g*_ distributions of the different time intervals for both simulations (Figure 5), which suggests convergent sampling has been achieved for both systems.

**Figure 5:**
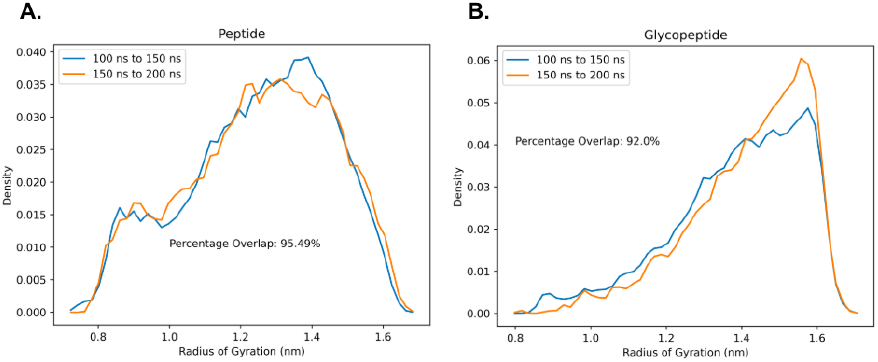
Probability distributions of *R*_*g*_ within the time intervals 100-150 ns and 150-200 ns of REMD simulation at 300 K for the 2 tandem repeat model peptides: A) without glycosylation and B) with O-linked GalNAc-Gal glycosylation. In both cases, there is good overlap between the *R*_*g*_ probability distributions within the two independent time intervals, indicating that the sampling has converged.

### Stability of the PPII Conformation

To explore the secondary structural landscape of the lubricin model peptide and how this landscape is influenced by lubricin’s O-linked GalNAc-Gal glycosylations, REMD simulations were used to construct the Ramachandran-like free energy surfaces at 300 K for both the non-glycosylated and glycosylated versions of the model peptide (Figure 6). For both systems, the global free energy minimum lies within the PPII helical conformation region, demonstrating the stability of the PPII helical conformation regardless of the presence of glycosylations. The PPII helical conformation stability was maintained with glycosylations, whereas right and left helical conformations were diminished. This suggests the most preferred conformation adopted by our model peptides is the PPII helical conformation, whose formation is likely driven, in large part, by the peptides’ sequence identity.

**Figure 6:**
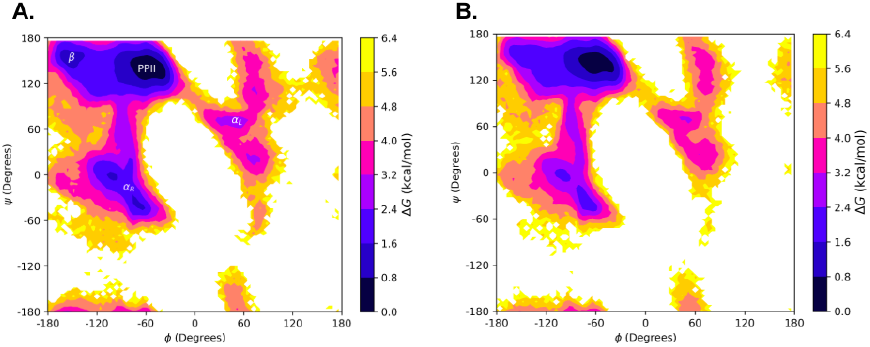
Ramachandran-like free energy surfaces at 300 K of the model lubricin peptide: A) without any glycosylations and B) with O-linked GalNAc-Gal glycosylations on relevant threonine residues (every threonine except the third one). The regions of the (*ϕ, ψ*) space corresponding to the polyproline type II conformation (PPII), beta conformation (*β*), right-handed alpha-helical conformation (*α*_*R*_), and left-handed alpha-helical conformation (*α*_*L*_) are labeled in A).

### Effect of O-linked Glycosylations on Threonines in the Model Peptides

Although the free energy surfaces for the lubricin model peptide with and without glycosylations are similar, slight differences between them may indicate that the glycosylations still play a role in facilitating the formation of the PPII helical conformation. Generally, regions outside of the PPII region tend to have higher relative free energies for the glycopeptide. For example, the relative free energy of the right-handed *α*-helical region is ∼0.8-2.4 kcal/mol in the non-glycosylated peptide, but ∼1.6-3.2 kcal/mol in the glycopeptide. A free energy local minimum of ∼0.8-1.6 kcal/mol is also present in a small area of the *β* region for the non-glycosylated peptide, but not the glycopeptide. The peripherals of each major secondary structural region, such as the transitional area between the PPII region and left-handed *α*-helical region, also tend to possess slightly higher relative free energies in the glycopeptide than in the non-glycosylated peptide. Hence, the presence of the O-linked GalNAc-Gal glycosylations could possibly play a role in promoting the structural order of the PPII helical conformation by slightly destabilizing conformations outside of the PPII region. However, examining these free energy surfaces alone is not sufficient to determine the exact effect of the glycosylations.

In order to obtain a more detailed picture of how the O-linked glycosylations affect the structure of the model peptide, we calculated the populations of each residue in both the glycosylated and non-glycosylated peptides for different secondary structural regions of the (*ϕ, ψ*)space using data from the 300 K replica of our REMD simulations. These regions are classified, according to the literature, as *α*_*R*_: −160^°^ *< ϕ <*−20^°^ and −120^°^ *< ψ <* 50^°^; *β*: −180^°^ *< ϕ <* −90^°^ and 50^°^ *< ψ <* 240^°^; PPII: −90^°^ *< ϕ <* 20^°^ and 50^°^ *< ψ <* 240^°^; *α*_*L*_: 30^°^ *< ϕ <* 100^°^ and 0^°^ *< ψ <* 80^°^; and coil for all other regions (42, 43). Only the five threonine residues possessed a practically and statistically significant difference (*α* = 0.05) in PPII population between the glycosylated and non-glycosylated peptides (Figure 7). Compared to the non-glycosylated peptide, all threonine residues in the glycopeptide had larger populations in the PPII region and almost zero population in the *α*_*R*_ region. We speculate that such a drastic decrease in *α*_*R*_ populations in the presence of glycosylations is due to the compactness of the *α*_*R*_ conformation, which may introduce unfavorable steric interactions between the bulky glycan chains and the threonine residues. This is consistent with previous observations in the literature on the structural effects of asparagine-linked chitobiose disaccharide glycosylations, which have been reported to destabilize the *α*_*R*_ conformation due to its steric encumbrance (44). As minimizing steric interactions is one of the main drivers of the PPII conformation, the higher relative stability of the PPII conformation in the glycopeptide may be attributed to relief of the steric interactions between lubricin’s glycosylations and the threonine atoms. It is also interesting to note that, despite not being directly linked to a glycan, the third threonine in the model peptide also possessed higher PPII populations and significantly lower *α*_*R*_ populations in the glycopeptide, possibly indicating cooperativity with the neighboring glycosylated threonine.

**Figure 7:**
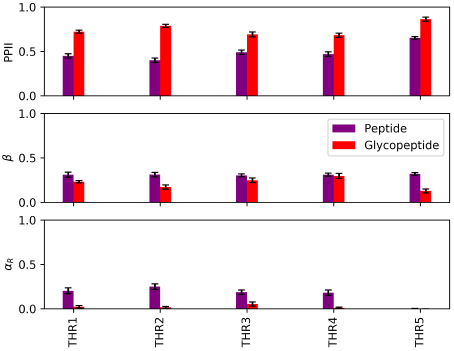
Secondary structure populations of the threonine residues in the model lubricin peptide (no glycosylations) and glycopeptide at 300K. Error bars represent 95% confidence intervals. The *α*_*L*_ and coil populations were negligible for both systems and are thus not included

### REMD Simulations of a Threonine Tetrapeptide

An important question to confirm is whether the O-linked GalNAc-Gal glycosylations increase threonine’s intrinsic propensity to adopt the PPII conformation, irrespective of the sequence identity of the surrounding amino acids. If this is true, then our results for the 2-tandem repeat lubricin model peptide can be more robustly generalized to the entire mucin domain, in which deviations in the sequence identity of the tandem repeats can occasionally occur. To address this question, REMD simulations were performed for a 4-residue TTTT peptide with and without O-linked GalNAc-Gal glycosylations on every threonine. Computational details for these simulations are essentially the same as those for the REMD simulations for the lubricin model peptide, except only 32 replicas with temperatures distributed in a geometric progression between 300 K and 452.32 K were used instead of 64 replicas. The simulations are well converged, as highlighted by the high overlap (*>* 90%) between the *R*_*g*_ distributions of two independent time intervals for both the glycosylated and non-glycosylated threonine tetrapeptide (Figure 8).

**Figure 8:**
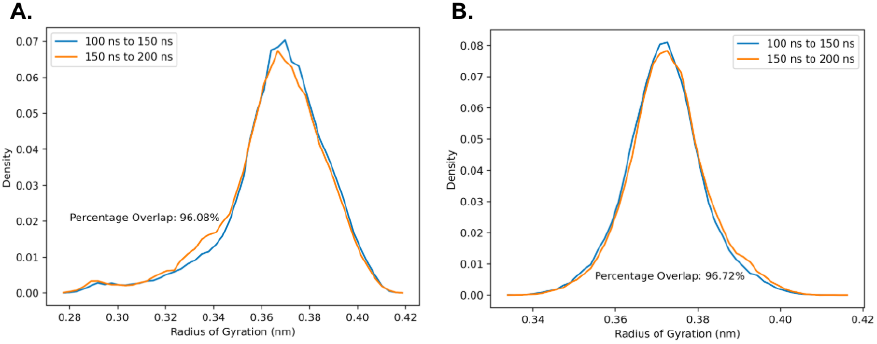
Probability distributions of *R*_*g*_ within the time intervals 100-150 ns and 150-200 ns of REMD simulations performed at 300 K for the 2 tandem repeat model peptides: A) Without glycosylation and B) With O-linked GalNAc-Gal glycosylation. In both cases, there is good overlap between the *R*_*g*_ probability distributions within the two independent time intervals, indicating that the sampling has converged.

Indeed, the presence of glycosylations resulted in drastic changes in the Ramachandran-like free energy surface of the threonine tetrapeptide (Figure 9). The addition of glycosylations essentially locks the threonine tetrapeptide in the PPII region, while the non-glycosylated threonine tetrapeptide can access other regions of the (*ϕ, ψ*) space much more freely. Ultimately, this indicates that the O-linked GalNAc-Gal increases threonine’s intrinsic propensity for adopting the PPII conformation, suggesting that the effects of the glycosylations on threonine would likely be similar regardless of the identity of neighboring residues.

**Figure 9:**
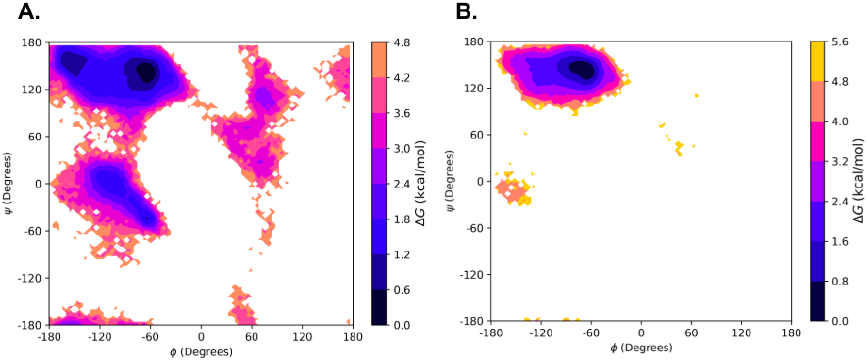
Ramachandran-like free energy surfaces at 300 K of threonine tetrapeptide: A) without glycosylations and B) with O-linked GalNAc-Gal glycosylations.

### Experimental Signatures of Polyproline Character in the PRG4 Mucin Domain: Circular Dichroism Spectra

Initial fractionation of rhPRG4 on a Sephacryl S500 HR column revealed two high molecular weight peaks, likely corresponding to oligomeric forms of the protein, dimer enriched (solid arrow) and dimer deficient (dashed arrow) (Figure 10A). Following a 1-hour digestion with Cathepsin G, these high molecular weight species were no longer present (Figure 10B), suggesting effective proteolysis and a higher concentration of lower molecular weight species. SDS-PAGE analysis of the second peak fractions from these chromatograms, with and without Cathepsin G digestion, revealed a major protein band at approximately 460 kDa, consistent with the expected size of monomeric rhPRG4 (Figure 10C). This band was also observed in the control sample, in which monomer, dimer and high-order oligomeric structures are seen as three distinct bands. Notably, in the Cathepsin G-digested sample, dimeric and high molecular weight species were absent, while they remained present in undigested conditions, further confirming that proteolytic digestion monomerized the protein. Western blot analysis using pAb1751 to probe against an epitope adjacent to the C-terminal subtilisin cleavage site (20) (Figure 2) shows these same bands as seen with SDS-PAGE, but also revealed a band at 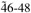 kDa (dotted arrow) (Figure 10D), which likely contains this epitope. The undigested rhPRG4 maintains a well-defined secondary structure with proportions of regular helices and turns. However, upon digestion, the secondary structure transitions to include regions that are mostly parallel beta-sheets and disordered regions (Figure 10E). This shift is evident from both the CD spectra and the BeStSel analysis, which collectively highlight the structural changes induced by digestion. The appearance of defined PPII content in the digested sample is further corroborated by the spectral data and reported structural analysis of anti-freeze glycoproteins (AFGPs), which form a triple helix of repeating Ala-Ala-Thr where the Thr residues are post-translationally modified with O-linked GalNAc-Gal (22).

**Figure 10:**
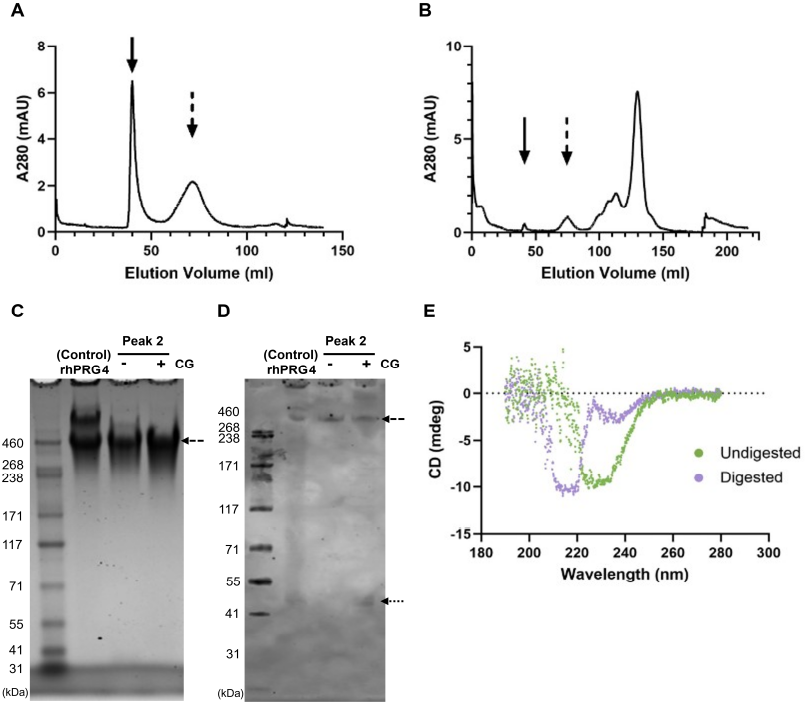
Cathepsin G digestion of rhPRG4 (lubricin) to partially isolate the mucin domain. A) Fractionation of rhPRG4 over Sephacryl S1000 in the presence of 1M NaCl in PBS at pH 7.4. B) Fractionation of the Cathepsin G digestate of rhPRG4 over Sephacryl S1000 showing digestion of the two principal high MW peaks (solid and dashed arrows). C) The partly digested second peak is detectable on SDS PAGE as a monomerized band with an apparent MW of 460 kDa (dashed arrow). Larger MW dimer and other larger MW forms are undetected. D) Western blotting of the same samples reveals immunopositivity with pAb1751 directed at the proximal side of the C-terminal subtilisin cleavage site observed by bands at 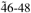 kDa (dotted arrow). E) Comparison of CD spectra of Cathepsin G digested and undigested rhPRG4 reveals spectral signatures consistent with polyproline II helix in the digested state which likely removed the C-terminal globular domain.

## DISCUSSION

Overall, our results provide novel insight into the dominant secondary structural element present within the PRG4 mucin domain. Although only a small subsection of the mucin domain could be simulated, our results for the lubricin model peptides should be generalizable to the rest of the mucin domain due to its repetitive nature. Additionally, prolines and threonines are the most common amino acids in the mucin domain, followed by lysines, alanines, and glutamates. Since proteins containing sequences of prolines, lysines, and alanines have all been shown in the literature to possess relatively high propensities for the PPII conformation (19), our observation that GalNAc-Gal glycoslyations increase threonine’s natural propensity for the PPII conformation means that the vast majority of amino acids in lubricin’s mucin domain highly prefer the PPII conformation, which further indicates the likely dominance of PPII helices within the mucin domain of PRG4.

Utilizing Cathepsin G to degrade the N- and C-termini with a limited digestion was supported by prior work by Karlsson *et al*. (39), who used the enzyme in an exhaustive digestion. Cathepsin G is known to cleave on the C-terminal side of the Lys and Arg residues that are well represented in the PRG4 primary structure in a trypsin-like manner (45). However, Cathepsin G is also known to display chymotrypsin-like activity, particularly around Phe residues. The motif of Pro-Phe appears to facilitate the most rapid proteolysis (45). This motif is located near the C-terminal end of the XX-TT-XX secondary mucin domain at Pro^970^-Phe^971^ within exon 6 of PRG4 (GenBank, NM_005807.6). The digestate contained a 46-48 kDa band observed on Western blot which is near the predicted molecular weight of 48.9 kDa encompassing Phe^971^ to Pro^1404^. This formed the basis for the selection of Cathepsin G in removing the N- and C-terminii since only one other aromatic residue is present in the PRG4 mucin domain at Phe^231^ and is not adjacent to a Pro residue. The digestion was brief, which likely preferentially spared sequences containing Lys and Arg. These digestion conditions exposed the main KEPAPTTP mucin domain, enabling the CD spectroscopy.

The mucin domain of PRG4 shares some structural similarities with an *α*-helix in that the characteristic heptameric repeating sequence abcdefg results in non-polar residues at positions “a” and “d” sitting near each other on the helical cylinder surface. These islets of hydrophobicity drive intermolecular attraction to other *α*-helices in an aqueous environment, resulting in the classical coiled-coil structure that drives dimerization and fibril formation. By comparison, the mucin domain of PRG4 is also characterized by a heptameric repeating degenerate sequence of KEPAPTTP (1, 3) in the conformation of a polyproline II secondary structure. This is topographically shaped like a prism and results in outward-facing isotactic GalNAc-Gal-NeuAc side chains, which are O-linked to threonines. The net result is a rigid structure, which at an extended contour length, will fold back upon itself and minimize exposure of hydrophobic alanines. The result is also an intra-molecular coiled-coil structure, which has been confirmed by surface force apparatus studies (4) in which PRG4 was observed to have a contour length of 220 +/-30 nm.

If our computational results are experimentally verified, then more work is required to link the mucin domain’s polyproline helical structures to other beneficial properties of lubricin, such as its ability to prevent gout by inhibiting nucleation of uric acid crystals (7). Perhaps, it may be possible to simulate a model lubricin peptide in the presence of uric acid crystals and observe how the model peptide interacts with uric acid’s tri-clinic crystalline structure, which is responsible for the needle-shaped crystals that drive inflammation (7, 46). This type of simulation would complement previous work conducted by Mochizuki *et al*., who simulated an antifreeze glycoprotein in the presence of ice crystals to determine the mechanism behind its antifreeze properties (22). Nonetheless, the structural insights obtained in the present work on lubricin’s mucin domain mark a step forward towards uncovering a full understanding of lubricin’s mechanism of action.

## CONCLUSIONS

REMD simulations of lubricin model peptides composed of two mucin domain tandem repeats have demonstrated the stability of PPII helices within lubricin’s central mucin domain. The threonine-linked GalNAc-Gal glycosylations within the mucin domain facilitate the adoption of the PPII conformation by destabilizing other secondary structural conformations accessible to the threonines, especially the right-handed *α*-helical conformation. If the PPII helical conformation is adopted, then a bottle brush polymer structure may emerge, which matches previous observations in the literature of lubricin’s end-grafted polymer-like properties, important in its surface-protecting, anti-adhesive, and lubrication activities (4, 5, 10).

## DATA AVAILABILITY

The datasets, code, and scripts that underlie the simulations discussed in this study may be found in our manuscript GitHub Repository (47).

## AUTHOR CONTRIBUTIONS

B.F., K.E., G.J., and B.R. conceived of this project. B.F. conducted all computational modeling, while A.M. and M.N. performed the CD spectroscopy. T.S. provided critical reagents. B.F., A.M., F.K., and B.R. wrote the manuscript. All authors edited the manuscript.

## ACKNOWLEDGMENTS

The authors would like to acknowledge insightful conversations with Gabriel Monteiro da Silva. B.F. is gracious for support from the Brown University SPRINT/UTRA and Research at Brown programs. G.J. acknowledges support from Brown Physicians Inc. G.J. and K.E. were funded by NIH R01 AR067748. B.R. was funded by NSF CTMC CAREER Award 2046744. The computational component of this research was conducted using computational resources and services at the Center for Computation and Visualization, Brown University.

## SUPPLEMENTARY MATERIAL

An online supplement to this article can be found by visiting BJ Online at http://www.biophysj.org.

